# Mis-perception of motion in depth originates from an incomplete transformation of retinal signals

**DOI:** 10.1101/443218

**Authors:** T. Scott Murdison, Guillaume Leclercq, Philippe Lefèvre, Gunnar Blohm

## Abstract

Depth perception requires the use of an internal model of the eye-head geometry to infer distance from binocular retinal images and extraretinal 3D eye-head information, particularly ocular vergence. Similarly for motion in depth perception, gaze angle is required to correctly interpret the spatial direction of motion from retinal images; however, it is unknown whether the brain can make adequate use of extraretinal version and vergence information to correctly interpret binocular retinal motion for spatial motion in depth perception. Here, we tested this by asking participants to reproduce the perceived spatial trajectory of an isolated point stimulus moving on different horizontal-depth paths either peri-foveally or peripherally while participants’ gaze was oriented at different vergence and version angles. We found large systematic errors in the perceived motion trajectory that reflected an intermediate reference frame between a purely retinal interpretation of binocular retinal motion (ignoring vergence and version) and the spatially correct motion. A simple geometric model could capture the behavior well, revealing that participants tended to underestimate their version by as much as 17%, overestimate their vergence by as much as 22%, and underestimate the overall change in retinal disparity by as much as 64%. Since such large perceptual errors are not observed in everyday viewing, we suggest that other monocular and/or contextual cues are required for accurate real-world motion in depth perception.

## Introduction

Stereoscopic vision is crucial for perceiving and acting on objects moving around us in three-dimensional (3D) space. Consider a batter in baseball: to accurately swing at an approaching pitch, the visuomotor system must first estimate the 3D spatial motion of the ball in space from two 2D retinal projections (Batista, Buneo, Snyder, & Andersen, 1999; Blohm & Crawford, 2007; Blohm, Khan, Ren, Schreiber, & Crawford, 2008; Chang, Papadimitriou, & Snyder, 2009). That means the brain has the difficult task of assigning coordinating points on each retina to the moving object and using an internal model of the eye-head geometry to accurately compute its 3D egocentric distance (Blohm et al., 2008). However, exactly which signals are used to extract motion-in-depth from binocular images is unclear.

Part of the confusion comes from an overabundance of available depth cues. Motion-in-depth cues can arise from both retinal and extraretinal sources and can be monocular or binocular. Monocular cues include retinal image features (e.g., shading, texture, defocus blur, perspective, optical expansion, kinetic depth cues, motion parallax, etc.) (Guan & Banks, 2016; Held, Cooper, & Banks, 2012; Zannoli, Love, Narain, & Banks, 2016; Zannoli & Mamassian, 2011), and ocular accommodation (Guan & Banks, 2016; M Mon-Williams & Tresilian, 2000). Binocular cues include retinal disparity, inter-ocular velocity differences, ocular vergence (Mark Mon-Williams & Tresilian, 1999; M Mon-Williams, Tresilian, & Roberts, 2000) and version angles (Backus, Banks, Van Ee, & Crowell, 1999; Banks & Backus, 1998). Ultimately, however, because retinal disparity varies non-uniformly with 3D eye-in-head orientation (Blohm et al., 2008), retinal signals alone are insufficient to estimate motion-in-depth; rather, the visual system must account for the full 3D geometry of the eye and head (Blohm et al., 2008). Indeed, Blohm et al. (2008)demonstrated that the visual system accounts for 3D eye-in-head orientation to accurately reach to static objects in depth, but how this finding extends to *moving* objects in depth is unclear. Here, we attempt to answer this question by asking participants to reconstruct motion-in-depth trajectories from only binocular depth cues across various horizontal vergence and version angles.

Another open question is how motion-in-depth perception depends on retinal eccentricity. Although the magnitude of binocular disparity increases with retinal eccentricity (Blohm et al., 2008), many of the observed disparity-selective cortical cells are tuned for small-magnitude disparities (DeAngelis & Uka, 2003), hinting that binocular signals may play a large role for depth perception near the fovea but not in the periphery. Convincing work from Held et al. (2012)found that position-in-depth is extracted in a complementary way: using mostly binocular disparity signals at the fovea and using mostly defocus blur in the periphery. Whether motion-in-depth estimates are similarly eccentricity-dependent, however, is unclear.

In this study, we asked participants to reproduce the perceived horizontal depth spatial trajectory of an isolated point stimulus observed either foveally or peripherally under different vergence and version angles. We found large systematic errors in the perceived motion trajectory that seemed to reflect an intermediate reference frame between purely retinal and spatial coordinates. A simple geometric model could capture the behavior well, revealing that participants tended to underestimate their version, overestimate their vergence, and underestimate the overall change in retinal disparity. These findings suggest that real-world motion-in-depth estimation is an eccentricity-dependent process that relies heavily on the use of monocular and/or contextual cues.

## Materials and methods

### Participants

In total, 12 participants (age 22-35 years, 9 male) were recruited for two experiments after informed consent was obtained. 11 of 12 participants were right-handed and all participants were naïve as to the purpose of the experiment. All participants had normal or corrected-to-normal vision and did not have any known neurological, oculomotor, or visual disorders. We also evaluated participants’ stereoscopic vision using the following tests: Bagolini striated glasses test (passed by all participants), Worth’s four dot test (passed by all participants), and TNO stereo test (all but 2 participants could detect disparities ≤60 seconds of arc). All procedures were approved by the Queen’s University Ethics Committee in compliance with the Declaration of Helsinki.

### Experimental paradigm

We used a novel 3D motion paradigm to determine how motion-in-depth is perceived across different horizontal version and vergence angles in complete darkness. This paradigm is illustrated in Fig. 1. In panel A, we show the physical setup with the array of red light-emitting diodes (LEDs) representing possible fixation targets (FTs; filled red circle represents the sample trial’s illuminated FT) and the green LED (filled green circle) representing the motion target (MT), which was attached to the arm of a custom 3D gantry system (Sidac Automated Systems, North York, ON) that was positioned at the same elevation as the eyes and moved within the horizontal depth (*x-y*) plane. At the end of target motion, participants were instructed to reconstruct the motion of this target using a stylus on the touchscreen in front of them. On each trial, the FT was reflected through a mirror oriented at 45° and positioned at the level of the eyes, such that the participant perceived the FT as located in the same horizontal depth plane as the MT. Other key elements in the physical setup included a stationary Chronos C-ETD 3D video-based eye tracker (Chronos Vision, Berlin, Germany) with an attached bite-bar for head stabilization to ensure stable fixation on the FT during target motion. This physical arrangement allowed us to present FTs in the MT plane while avoiding physical collisions (panel B) with FTs positioned at nine different locations (corresponding to three horizontal version angles, −30°, 0° and 30°, and three vergence angles, 3°, 4.8° and 8.8°) and 18 different motion trajectories (six orientations spaced equally from 0° to 180°, with three possible curvatures) purely in the horizontal depth plane.

**Figure 1:**
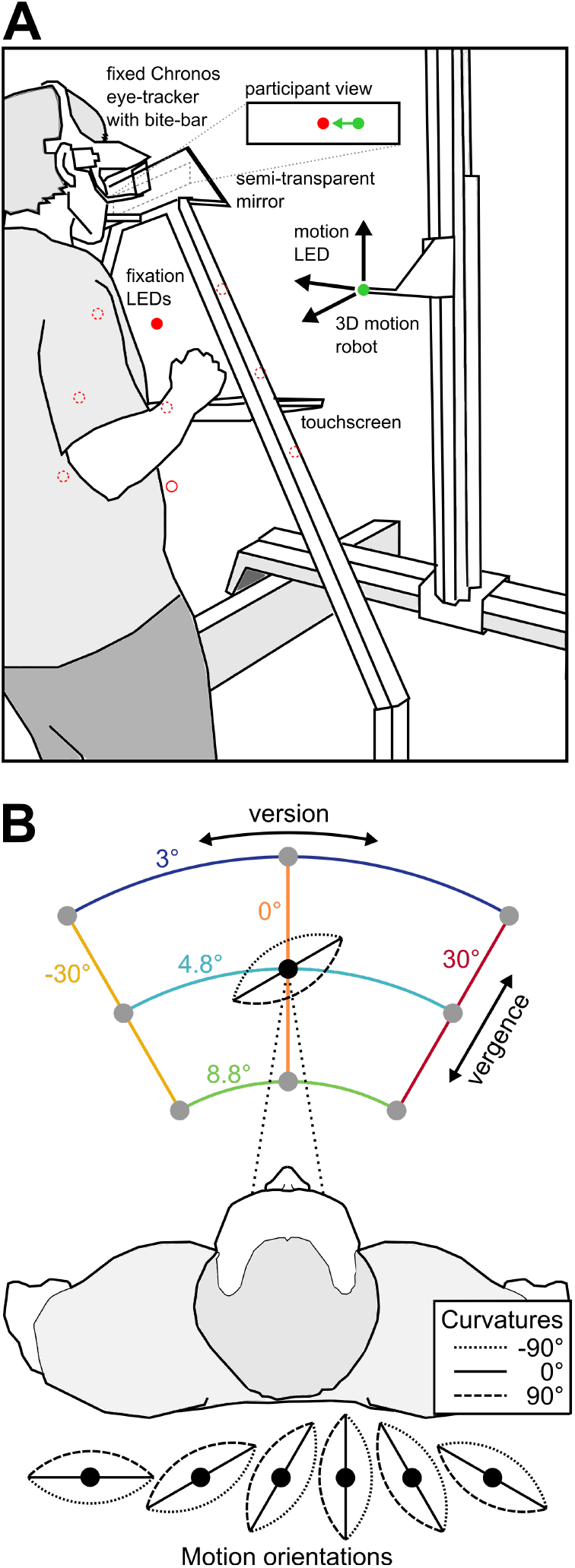
Apparatus and virtual setup. **A** Experimental apparatus, including 3D motion robot with attached MT LED (green), frontoparallel arc-array of 9 FT LEDs (red), 45° oriented semi-transparent mirror, fixed Chronos eye tracker, and touchscreen. For a given trial, one of the FT LEDs is illuminated and reflected at eye-level using the semitransparent mirror. Meanwhile, the motion robot moves the MT LED in the horizontal depth plane also at eye level, creating the participant view shown in the inset. **B** Virtual setup created by the experimental apparatus and tested motion trajectories, with 6 orientations (30° steps from 0° to 150°) and 3 curvatures (−90°, 0° and 90°).

### Procedure

Participants knelt, supported by the custom apparatus, in complete darkness. Each trial was defined by three phases: (1) fixation, (2) motion observation and (3) reporting. During the fixation phase (0 ms –1500 ms), participants fixated a randomly selected, illuminated FT from the array of nine LEDs. During the motion observation phase (1500 ms –3200 ms), participants maintained fixation on the FT while the MT was displaced by the robot. That MT displacement either occurred in the immediate space around the FT (foveal condition) or around the central (non-illuminated) LED while the participant maintained fixation on the FT (peripheral condition). Participants were asked to memorize its trajectory in the *x-y* plane. During the reporting phase (3200 ms – trial end), participants were asked to remove their head from the bite-bar and trace the perceived spatial trajectory using a stylus on a touchscreen, illuminated using a single bright LED for this trial phase only. The light remained on until a response was recorded, and participants were free to restart their trace at any time. They touched the lower right corner of the screen in order to end the current trial, triggering the start of the next trial.

### Trial selection

We recorded a total of (9 fixation targets * 3 curvatures * 6 orientations * 2 motion location conditions =) 324 trials for each participant (324 trials * 13 participants = 4212 total trials). Each trial type was randomly interleaved throughout 10 blocks (per participant) but the order was the same across all participants. This allowed us to pool the responses together across conditions and participants for graphical purposes, as there were no within-participant trial repetitions (model fits were performed on individual trajectories). Upon offline analysis, we discovered that one participant consistently failed to perform the reconstruction portion of the task as instructed: the participant drew the motion backwards and we therefore excluded his data from the analysis, leaving 12 participants (3888 total trials). Of these trials, we examined recorded eye movement data and removed trials containing eye movements or blinks during the motion phase of each trial, leaving 3869 valid trials for analysis.

### 3D binocular kinematic model

We developed a 3D model of the binocular retina-eye-head geometry to predict how behavioral motion reconstructions might vary across version and vergence angles (Blohm et al., 2008). This model consisted of three primary stages: retinal motion encoding, inverse modeling and spatial motion decoding. First, we computed the binocular retinal projections of the motion stimulus, given the current eye and head orientations (retinal motion encoding stage). Second, we moved the eyes together to the inverse estimates of version and vergence angles during the encoding of retinal motion (inverse modeling stage). We then back-projected the retinal coordinates into space and computed the 3D location of the rays’ intersection, representing the decoded depth (spatial decoding stage). Although in reality we computed all three stages of this model to obtain our trajectory estimates, we simplify the graphical representation of this model in Figure 2: focusing on the inverse modeling stage, where we varied the contributions of extraretinal signals.

**Figure 2:**
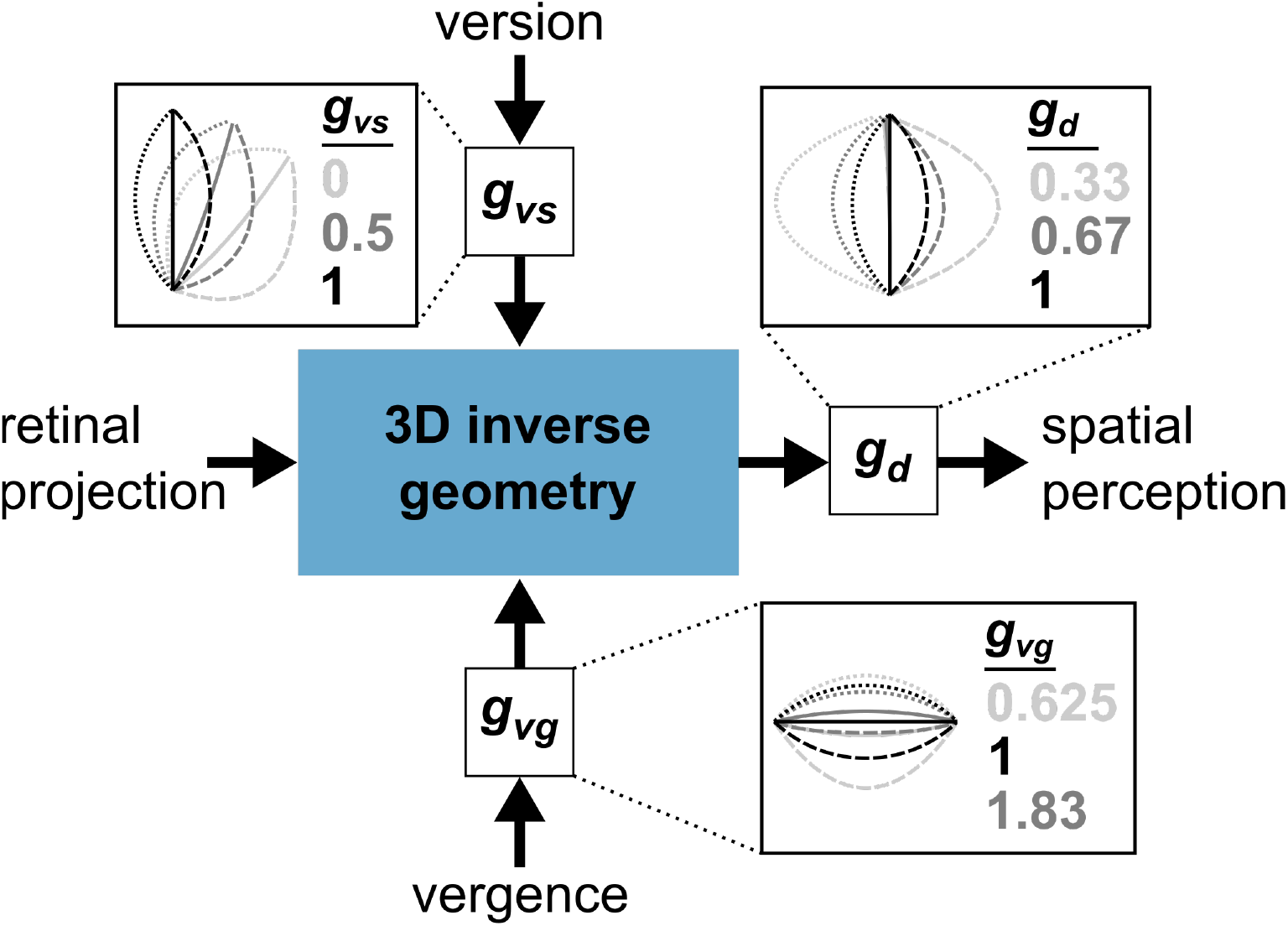
Inverse modeling stage of the 3D binocular kinematic model to generate retinal and partial model predictions. Insets show individual parameter effects on reconstructed traces. The effects of version gain are shown for a fixation version angle of 30deg; the effects of vergence gain are shown for a fixation vergence of 4.8deg; the effects of depth gain are shown for trajectories with an orientation of 90deg.

This modeling framework allowed us to describe the reconstructed trajectories by varying the contributions of version (version gain, *g_vs_*) and vergence (vergence gain, *g_vg_*) to the inverse model, and motion purely in depth (depth gain, *g_d_*). Each parameter accounted for a different aspect of the trajectory (shown in Figure 2 insets). To produce the retinal prediction, we set the version gain to 0 and used a constant vergence gain of 1. Importantly, this retinal prediction arbitrarily assumes that vergence is 100% accounted for. Note that, because our model computes the spatial intersection of the binocular back-projections, vergence gains had to be greater than 0 (otherwise the back-projections would be parallel).

For each participant, we initialized the parameters for the reconstructed trajectories using a brute-force, 8000 point least-squares method over the full plausible range of parameters (20 linearly spaced values for each parameter). This was followed by a 512 point least-squares fine fitting method within a +/− 10% range for *g_vs_, g_vg_* and *g_d_* around the initialized parameters (8 linearly spaced values for each parameter). We performed this exact optimization procedure separately for each vergence angle to avoid confounding vergence effects. In total, we computed the fits of (3*(8000+512) =) 25,536 total parameter combinations. This optimization provided parameter estimates that consistently accounted for behavioral variability, with each participant’s R-squared values >0.93 in both motion conditions.

### Statistical analyses

Group-level statistical tests primarily consisted of two-tailed Student t-tests. We also performed paired t-tests when appropriate for comparing parameters across conditions. The rest of the statistical treatment of the data consisted primarily of computing correlation coefficients and regression analyses.

## Results

We sought to determine how visual perception accounts for binocular eye orientation when reconstructing motion in depth. To do this, we designed a novel paradigm in which participants reconstructed motion of an LED in the horizontal depth plane presented either foveally or peripherally on the retina, while fixated in one of nine randomly selected version and vergence orientations. After observing the motion, participants generated this reconstruction using a touchscreen positioned in the coronal plane directly in front of them. We then analyzed these reconstructed trajectories to determine how they varied across eye orientation and motion condition. To generate model predictions for the reconstructed signals across changes in version and vergence angles, we developed a 3D model of the binocular eye-head geometry (Figure 2, see *Methods* for model details). This model allowed us to characterize the eye orientation signals accounted for by the perceptual system.

Reconstructed trajectories deviated from both the spatial (physical) and retinal (see Methods) predictions for both foveal and peripheral motion across all vergence angles. These trajectories are shown alongside their predictions for three representative motion orientations in Figure 3, averaged across all participants and vergence angles for foveal (top row) and peripheral motion (bottom row). These comparisons revealed both an angular displacement between the trajectories during nonzero version as well as a compression of the behavioral traces in the depth dimension across motion orientations; however, these patterns were not consistent for both motion conditions, as only the compression effect was obvious in the peripheral case.

**Figure 3:**
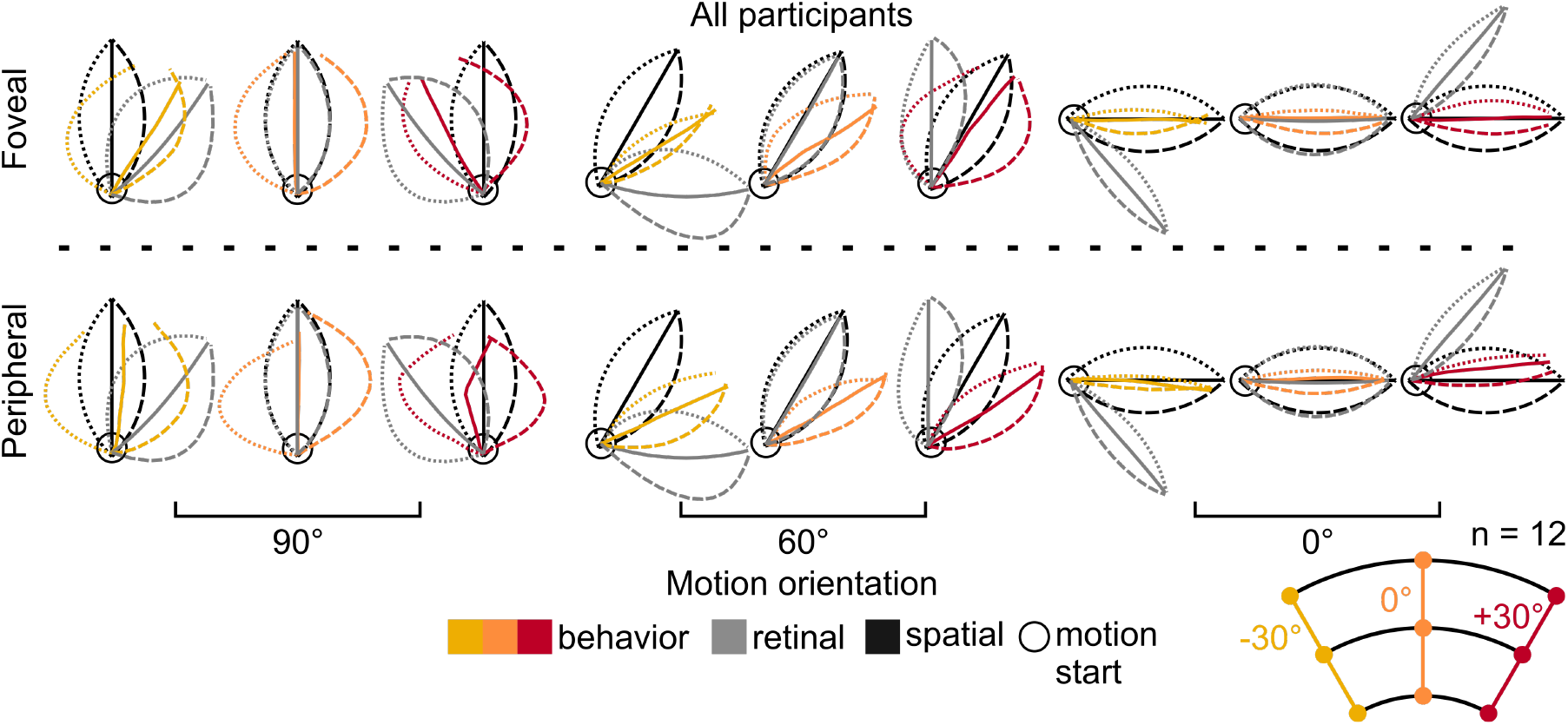
**Average across-participant foveal and peripheral reconstructed motion**, compared with spatial and retinal predictions. Note that reconstructed traces were normalized in amplitude to the spatial and retinal predictions.

The reconstructed trajectories matched neither the spatial nor retinal hypotheses, suggesting that the perceptual system only partially transformed the retinal MT trajectories into spatial coordinates. To capture the extent to which the perceptual system encoded binocular eye orientations and estimated motion purely in the depth dimension (i.e. when the MT was stationary on the retina), we used a two-step least-squares algorithm to optimize the ***g_vs_***, ***g_vg_*** and ***g_d_*** inverse model parameters for the behavioral trajectories (see *Methods* for detailed explanation of optimization algorithm). The results of this optimization are shown in Figure 4 at both the single participant level (panel A) and group level (panel B) for both the foveal and peripheral motion conditions.

**Figure 4:**
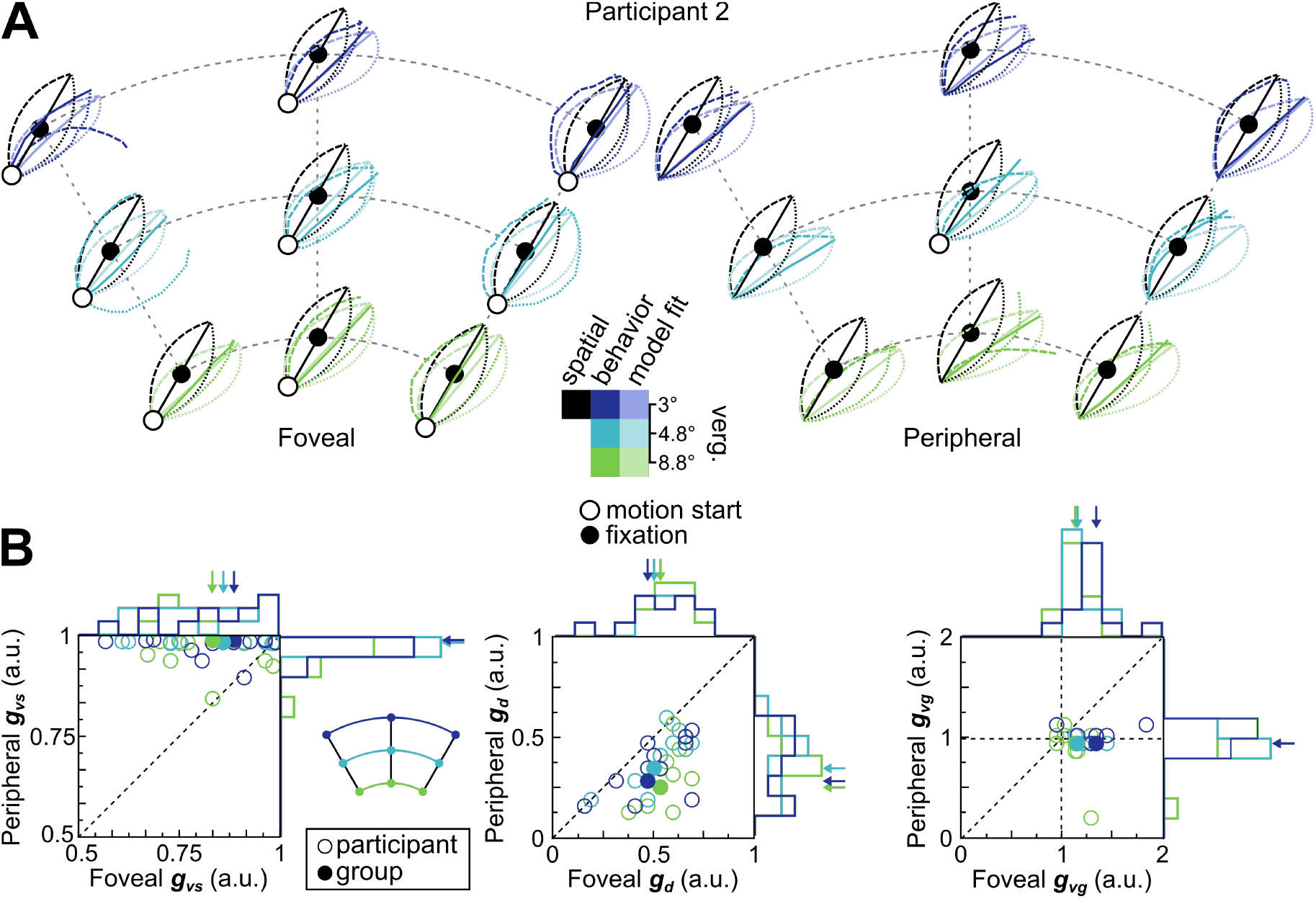
Results of model optimization. **A** Comparison of model outputs and spatial predictions with actual reconstructed trajectories after fitting version gain (***g_vs_***), depth gain (***g_d_***) and vergence gain (***g_vg_***) parameters separately for each vergence distance and for foveal (*left*) and peripheral (*right*) motion conditions, for single participant (#2). **B** Group-level scatter plots showing peripheral versus foveal motion parameter fits for version gain (***g_vs_***, *left*), depth gain (***g_d_***, *middle*) and vergence gain (***g_vg_***) (*right*). Open disks represent participant parameters and solid disks represent group-level parameters fit on all the data. Arrows above histograms represent group-level fit parameter locations along a given axis.

The parameters optimized for foveal and peripheral motion were distinct, suggesting that motion-in-depth perception varies with retinal eccentricity. For version gain, we found that participants accounted for 83% +/− 13% (mean +/− SD) of horizontal version during foveal motion, compared to 96% +/− 10% during peripheral motion (paired t-test: t(35) = −5.22, p < 0.01). Given that version compensation during foveal motion was incomplete, the apparent full compensation during peripheral motion could have been the result of the system using the retinal location of the stimulus as a cue for current horizontal eye orientation, effectively bypassing an explicit need for extraretinal signals. Next, we found that the foveal depth gains accounted for 54% +/− 13% of depth speed and was significantly greater than that for peripheral motion at 36% +/− 14% (paired t-test: t(35) = 8.70, p < 0.01), indicating that motion in depth was perceived to be faster when foveal. Finally, participants used a foveal vergence gain of 1.22 +/− 0.18. In contrast, participants used a significantly smaller (and more accurate) peripheral vergence gain of 0.98 +/− 0.15 (paired t-test: t(35) = 6.30, p < 0.01). These findings suggest that an underestimation of 3D eye orientation signals during the transformation from retinal to spatial coordinates is responsible for observed distortions to motion-in-depth perception.

Taken together, these three fit parameters allowed us to characterize the extent to which the transformation from retinal to spatial coordinates occurred for each participant. We computed a transformation index, *I_T_*, represented by equation (1):

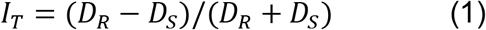

Where *D_R_* and *D_S_* are the Euclidian distances of each set of gain parameters from the retinal and spatial hypotheses, respectively. For example, a purely spatial set of gain parameters would be represented by [*g_vs_ g_vg_ g_d_*] = [1 1 1], corresponding to a *D_S_* = 0 and a D_R_ = 1; subsequently, *I_T_* = (1-0)/(1+0) = 1. By the same logic, for a purely retinal set of gains, *I_T_* = −1. We present the distributions of these gain parameters for each participant separated for foveal and peripheral motion, merged across vergence fits, in Figure 5. *I_T_* was significantly greater than 0 for both foveal (mean +/− SD: 0.48 +/− 0.12; t(35) = 24.4, p < 0.01) and peripheral motion (mean +/− SD: 0.37 +/− 0.17; t(35) = 13.3, p < 0.01), suggesting that, in both cases, reconstructed trajectories were intermediate, but more spatial than retinal.

**Figure 5:**
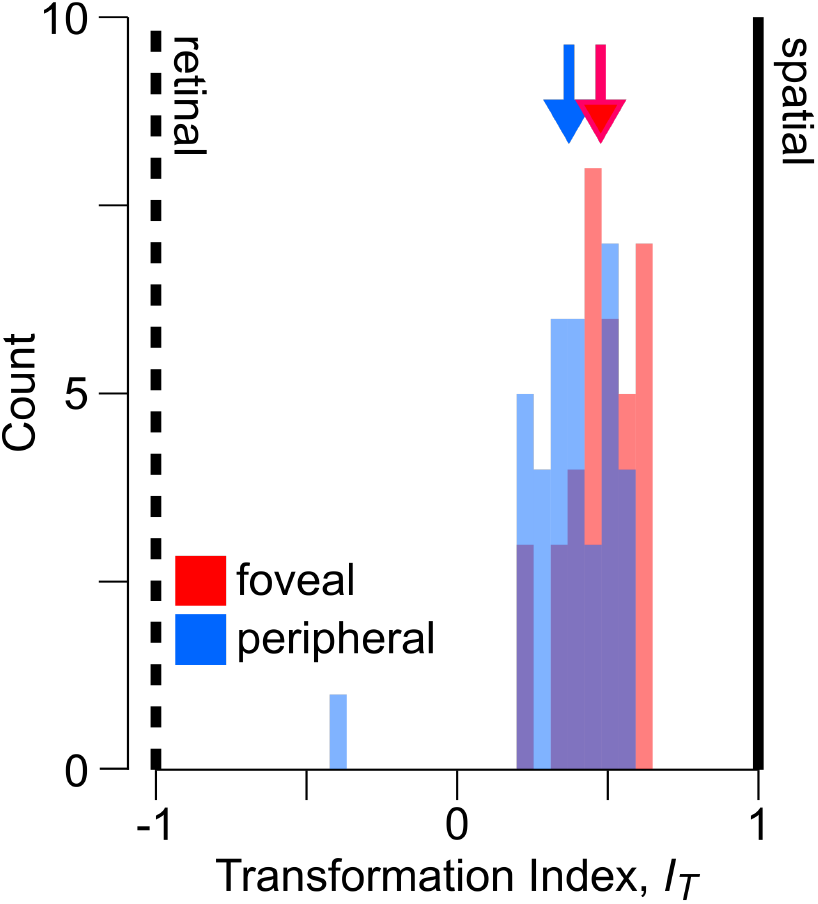
Transformation indices (*I_T_*) for foveal and peripheral motion. Also shown are the retinal (dashed) and spatial (solid) predictions, with means for foveal (red) and peripheral (blue) motion represented by color-matched arrows.

## Discussion

We asked participants to estimate the motion-in-depth of an isolated disparity stimulus and found large systematic errors that differed depending on viewing eccentricity. We found that a simple model of the 3D eye-in-head geometry that used inverse estimates of the ocular version angle, vergence angle and speed-in-depth could capture the reconstructed trajectories well. For foveal motion, a model that overestimated ocular vergence angle, underestimated ocular version and target speed fit the perceived trajectories. For natural viewing, this result suggests that additional monocular cues are necessary to accurately estimate foveal motion in depth. Contrastingly, for peripheral motion, a model that accurately estimated eye orientation signals fit the perceived reconstructions, but this model also severely underestimated the speed in depth – more than during foveal motion. In this condition, binocular eye orientations may have been inferred using eccentricity-related monocular cues. Using a simple transformation index computed from the inverse model fits, we found that spatial misperceptions corresponded to a partial transformation of retinal motion into spatial coordinates, regardless of retinal eccentricity.

We found that the visual system cannot use binocular retinal signals alone to accurately estimate motion-in-depth at the fovea during fixation. This reliance on monocular retinal cues and/or contextual depth cues is understandable given their abundance in natural vision; however, absolute depth cannot be extracted from monocular cues alone. Of course this is rarely an issue for typical viewing when multiple relative depth cues are available. An additional reason why the visual system might rely on often geometrically inaccurate cues potentially comes from the idea that estimates of binocular eye orientation are unreliable (Blohm et al., 2008; McGuire & Sabes, 2009), and these estimates might become even more variable due to stochastic reference frame transformations (Alikhanian, Carvalho, & Blohm, 2015). The head-to-world-centered coordinate transformation required to extract depth from disparity could be stochastic (i.e., adding uncertainty to the final depth estimate) and the fovea’s high spatial acuity means that monocular motion cues could be quite reliable for stereopsis (e.g., Ponce and Born, 2008) This interpretation is consistent with various lines of evidence showing a propensity of the visuomotor system to optimally account for perceptual (Jessica K Burns & Nashed, 2011) and motor uncertainty (Jessica Katherine Burns & Blohm, 2010; Schlicht & Schrater, 2007; Sober & Sabes, 2003) resulting from stochastic reference frame transformations (Alikhanian et al., 2015), and is consistent with behavioral evidence from visuomotor updating work (Fiehler, Rösler, & Henriques, 2010; Henriques, Klier, Smith, Lowy, & Crawford, 1998; Medendorp, Goltz, Vilis, & Crawford, 2003; Murdison, Paré- Bingley, & Blohm, 2013). Finally, the tendency of disparity-tuned neurons to disproportionately prefer disparities <1 deg (DeAngelis & Uka, 2003) is another clue that motion-in-depth at the fovea is represented differently in the visual system than motion-in-depth in the periphery, where disparity magnitudes are much larger (Blohm et al., 2008).

The peripheral motion case presents an apparently paradoxical finding: eye-in-head orientation can be accurately estimated (likely using retinal eccentricity) while target speed-in-depth is significantly underestimated relative to both its spatial motion and its foveal motion. However, we provide *only* a disparity stimulus to the observer regardless of retinal location, and the relative contribution of disparity to depth perception decreases with eccentricity (Held et al., 2012). The observed percept of compressed motion in the periphery is therefore in line with the idea of a lower-weighted contribution of disparity cues (Held et al., 2012), while changes in defocus blur of the point stimulus were likely negligible. In agreement with this idea, some early psychophysical findings reveal that such a lateral compression could be due to greater relative uncertainty in the estimate of the depth motion component for motion in the periphery (Rokers et al., 2017, pre-print). Determining whether motion-in-depth perception is based on such a statistically optimal combination of disparity, retinal defocus blur and extraretinal cues therefore represents a potential extension of this work.

To isolate for horizontal disparity as the primary cue for depth perception, we removed any contribution of visuomotor feedback by restricting movements of the eyes and head. We determined the role of *static* eye orientation signals in interpreting a *dynamic, moving* stimulus, although in natural viewing our eyes and head are often moving as well. Both disparity and eye movements contribute to depth perception but the precise nature of these contributions, and how they might depend on one another, is unclear. For example, vergence angle corresponds to perceived depth during the kinetic depth effect (Ringach, Hawken, & Shapley, 1996), but artificially inducing disparity changes between correlated (and anti-correlated) random-dot stimuli can cause the eyes to rapidly converge (or diverge) without any perception of depth (Masson, Bussettini, & Miles, 1997). On the neural level, disparity is coded in V1 without a necessary perception of depth (Cumming & Parker, 1997). Psychophysics work has shown that vergence eye movements are beneficial for judging the relative depth of stimuli (Foley & Richards, 1972), but to our knowledge no one has investigated the extent to which these signals are used to solve the geometry for absolute depth.

In addition, by restricting the orientations of the eyes and head we removed feedback due to motion parallax and changes in vertical disparity. Importantly, providing such dynamic feedback has been shown to improve motion-in-depth perception in virtual reality (Fulvio & Rokers, 2017). Although vertical disparity naturally varies during normal ocular orienting, we designed our task to keep vertical disparity constant for a given gaze location. This manipulation not only removed vertical disparities due to changes in cyclovergence, but also vertical disparities due to changes in head orientation (Blohm et al., 2008). These natural changes in vertical disparity during eye and head movements likely serve as another informative dynamic cue for judging motion-in-depth under normal viewing contexts. For the above reasons, presenting participants with a dynamic, motion-tracked version of our task could therefore represent an important extension of this work.

From an evolutionary perspective, it is unclear why the visual system would underestimate binocular cues when estimating motion in depth with static gaze. Indeed, in an enriched visual environment there are often sufficient monocular cues available to the visual system to be able to judge relative depth. During everyday viewing in natural contexts, this is often the case; especially for self-generated motion in depth. On the other hand, our findings suggest that in some special cases without an enriched viewing context such a monocular strategy fails. To illustrate this point, consider two edge cases: juggling and firefly-catching. Expert jugglers learn to fixate the apex of the balls’ trajectory, presumably taking advantage of a learned internal model of the balls’ ballistic trajectory (resulting from manual motor commands) combined with various monocular motion cues to intercept each ball. Alternatively, consider the case of attempting to catch a firefly in darkness: fixating while attempting this is intuitively a bad idea because the flight of a firefly is largely unpredictable. Instead, to catch the fly, a better strategy might be to visually track its motion. Such a strategy would allow for the use of consistent visuomotor feedback, allowing the construction of a predictive model of the fly’s path. Thus, follow-up experiments investigating the interplay between (1) availability of monocular cues, (2) predictability of object physics and (3) facilitation from visuomotor learning would be informative of how our brain constructs motion-in-depth percepts.

## Conclusions

We quantified the extent to which visual perception accounts for the 3D geometry of the eyes and head when interpreting motion in depth under static viewing conditions. We found that participants underestimated 3D binocular eye orientations, leading to different spatial motion percepts for identical egocentric trajectories. To perceive and successfully navigate through the 3D world, our findings suggest that perception must supplement binocular disparity signals with binocular eye and head orientation estimates, monocular depth cues and dynamic visuomotor feedback. It remains to be seen, however, what the precise contributions and relative weightings of each of these cues might be.

## Acknowledgments

The authors want to thank ________. This work was supported by NSERC (Canada), CFI (Canada), the Botterell Fund (Queen’s University, Kingston, ON, Canada) and ORF (Canada). TSM was also supported by DAAD (Germany) and DFG (Germany).

